# Controlling for contaminants in low biomass 16S rRNA gene sequencing experiments

**DOI:** 10.1101/329854

**Authors:** Lisa Karstens, Mark Asquith, Sean Davin, Damien Fair, W. Thomas Gregory, Alan J. Wolfe, Jonathan Braun, Shannon McWeeney

## Abstract

**Background:** Microbial communities are commonly studied using culture-independent methods such as 16S rRNA gene sequencing. However, one challenge in accurately characterizing microbial communities is exogenous bacterial DNA contamination. This is particularly problematic for sites of low microbial biomass such as the urinary tract, placenta, and lower airway. Computational approaches have been proposed as a post-processing step to identify and remove potential contaminants, but their performance has not been independently evaluated.

To identify the impact of decreasing microbial biomass on polymicrobial 16S rRNA gene sequencing experiments, we used a serial dilution of a mock microbial community. We evaluated two computational approaches to identify and remove contaminants: 1) identifying sequences that have an inverse correlation with DNA concentration implemented in Decontam and 2) predicting the proportion of experimental sample arising from defined contaminant sources implemented in SourceTracker.

**Results:** As expected, the proportion of contaminant bacterial DNA increased with decreasing starting microbial biomass, with 79.12% of the most dilute sample arising from contaminant sequences. Inclusion of contaminant sequences in analyses leads to overinflated diversity estimates (up to 12 times greater than the expected values) and distorts microbiome composition. SourceTracker successfully removed over 98% of contaminants when the experimental environments are well defined. However, SourceTracker performed poorly when the experimental environment is unknown, failing to remove the majority of contaminants. Decontam successfully removed 74-91% of contaminants regardless of prior knowledge of the experimental environment.

**Conclusions:** Our study indicates that computational methods can reduce the amount of contaminants in 16S rRNA gene sequencing experiments. The appropriate computational approach for removing contaminant sequences from an experiment depends on the prior knowledge about the microbial environment under investigation and can be evaluated with a dilution series of a mock microbial community.

## Background

Advances in genomic sequencing have transformed our ability to identify and study microbes without depending on culture-based laboratory techniques. A common method used to study microbial communities is to sequence marker genes, such as the 16S rRNA gene. In this method, the bacterial DNA is extracted from a sample, amplified by PCR, and then sequenced. This technique is relatively inexpensive and easy to perform, and has increased our ability to understand the role of microbes in a variety of environments, including those of low microbial biomass such as the upper atmosphere[1], placenta [2], and urinary bladder[3–5].

The discovery of microbes in these unexpected environments is exciting and can revolutionize our understanding of these environments. However, a major challenge hindering our ability to accurately characterize microbial communities in these environments is bacterial DNA contamination from exogenous sources that are introduced during sample collection and processing. Contaminant bacteria introduced to the sample prior to PCR can dominate the composition of low microbial biomass samples[6–9], comprising over 80% of the sample in extreme cases[10]. Failure to account for contaminants may also affect biological conclusions drawn from studies, such as inflating alpha diversity metrics, distorting the abundance of true microbial members of the environment[10], and altering differences between clinical groups[7, 9, 11].

Currently, there is no standard method to minimize or control for contaminants in 16S rRNA gene sequencing experiments. During sample processing, procedures can be taken to minimize the amount of exogenous DNA introduced to the samples. These include pretreating reagents in an attempt to remove exogenous DNA[10, 12] and using DNA extraction kits designed specifically to minimize contamination. However, techniques involving pretreating reagents are challenging and may be ineffective when examining low microbial biomass samples[10].

Another approach is to process “no template” negative controls along with experimental samples through DNA extraction, PCR amplification, and sequencing steps. Even if no band is present on an agarose gel, these negative controls should be sequenced since band-negative samples have been reported to produce sequence reads[7]. While this is an effective strategy for identifying experiment-specific contaminants, there is little guidance for researchers regarding the appropriate use of this information. Removing all sequences found in the blank sample is not plausible since low levels of real sequences from the sequence run may be present in the blank control due to multiplexing artifacts[13, 14]. This leads to removing bacterial sequences that are actually biologically relevant; thus, this approach has been found to be too strict[10].

Several approaches to objectively remove contaminants after 16S rRNA gene sequencing have been suggested. One directly identifies and removes sequences that have been previously identified as contaminants in published databases or reference lists[7]. However, this is a generalized approach and might not accurately reflect the contamination present in the actual experiment. A second approach applies an abundance filter to remove all sequences that are below a defined threshold[15, 16]. However, this does not remove the abundant contaminants that are present in low microbial biomass samples and would also remove real rare sequences in the dataset. Furthermore, without the appropriate experimental controls, there is no guidance to determine the appropriate filter. A third approach is to remove sequences that are present in a negative control sample[17, 18]. However, this approach may be too harsh due to sample cross-contamination[14]. A fourth analytic approach is to identify bacterial reads that have an inverse correlation with bacterial DNA concentration after 16S rRNA library preparation[9, 15, 19]. This has been recently implemented in an open source R package called decontam[20]. A fifth method for identifying exogenous DNA contaminants uses a Bayesian approach implemented in SourceTracker[21] to predict the proportion of an experimental sample that arose from a defined contaminant source.

In this study, we use a dilution series of a mock microbial community to determine the success of current approaches to control for laboratory contaminants in studies involving low microbial biomass samples. Mock microbial communities are mixtures of known bacteria and are an important tool to identify biases in both laboratory and computational methods employed in microbial community analyses. We demonstrate that a dilution series of a mock microbial community is a valuable tool to evaluate optimal inputs and parameters for two bioinformatics approaches that have been proposed for contaminant identification from low microbial biomass samples.

## Methods

To identify the presence of exogenous bacterial DNA and its impact on low microbial biomass samples, we used 16S rRNA gene sequencing of a mock microbial community that had undergone eight rounds of serial three-fold dilutions. We chose a mock microbial community since most microbiome studies aim to identify mixtures of bacteria rather than pure bacterial isolates. In mock communities, the expected 16S rRNA gene sequences are known, thus any unexpected sequences identified in the analysis of the dilution series can be attributed to error. These errors can arise from sequencer errors (base miscalls or barcode switching), chimeric sequences, or laboratory contaminants.

### Microbial mock community dilution

ZymoBIOMIC mock community standards were used (Zymo Research). This mock community consisted of 8 bacterial species (*Pseudomonas aeruginosa, Escherichia coli, Salmonella enterica, Lactobacillus fermentum, Enterococcus faecalis, Staphylococcus aureus, Listeria monocytogenes, Bacillus subtilis*) and two fungal species (*Saccharomyces cerevisiae* and *Cryptococcus neoformans*). The mock community was diluted with microbial free water (Qiagen) in 8 rounds of a serial three-fold dilution prior to DNA extraction. A total of 50 μl of community standard (equivalent to ∽1.5×10^9^ total bacteria) was used as the highest microbial standard for DNA extraction (see Supplemental Table 1). Blank controls of microbial-free water also were used and subjected to all processing steps.

### DNA extraction and PCR amplification

A total of 50 μl of each serially diluted microbial standard was subject to DNA extraction using the cultured cell protocol supplied with the DNeasy Blood and Tissue Kit (Qiagen, Germany) per manufacturer’s instructions. DNA was eluted in a total volume of 50 μl. The extracted DNA was quantified and quality checked at A260 / A280 nm (Nanodrop, Thermo Fisher Scientific, USA) prior to amplification by polymerase chain reaction (PCR). Bacterial DNA was amplified by PCR using Golay barcoded primers, which target the V4 region of 16S rRNA genes (Caporaso et al., 2012). Template DNA was amplified in triplicate using the GoTaq Hot Start Polymerase kit (Promega, USA). One microliter of template DNA and 1 μL of a unique barcoded reverse primer were added to 48 μL of master mix containing 1x colorless reaction buffer, 1.5 mM MgCl_2_, 0.2 mM dNTPs, 0.2 mM forward primer, and 1.25 U of polymerase enzyme. The reaction volumes were placed in a thermocycler and run through the following conditions: 94°C for 3 min (initial denaturation), followed by 35 cycles of 94°C for 45 s (denaturation); 55°C, 40 s (annealing); 72°C, 1.5 min (extension); with a final extension at 72°C for 10 min.

### PCR product purification and sample pooling

Ten microliters of each product was used to verify amplification by gel electrophoresis on a 2% agarose gel. Replicates yielding visible bands at 382 bp were pooled together and purified following the QIAquick PCR Purification kit (Qiagen, Germany) provided protocol. Purified products were again quantified and quality checked at A260/A280 nm (Nanodrop, Thermo Fisher Scientific, USA). Products were diluted to 10 ng / μL and 5 μL of each sample were pooled together for sequencing on the Illumina MiSeq sequencer (Illumina, USA).

### Sequencing and sequence processing

Primers and sequence adapters were removed with the Illumina MiSeq Reporter (version 2.5). The sequences were further processed using scripts implemented in the R statistical computing environment with the DADA2[22] (v. 1.4.0) package (scripts are available on GitHub (https://github.com/lakarstens/ControllingContaminants16S)). Briefly, sequences were quality filtered, trimmed (forward reads at 240 nt, reverse reads to 160 nt) prior to inferring amplicon sequence variants (ASVs) with the DADA2 algorithm. ASVs, which group similar sequences together according to a model that considers sequence abundance and sequencing error, where chosen over traditional Operational Taxonomic Units (OTUs) since they have a finer resolution[23, 24]. Chimeric sequences were removed with the approach implemented in the DADA2 package. Taxonomy was assigned for each ASV to the genus level using the RDP naive Bayesian classifier[25] implemented in DADA2 with the SILVA database (version 123). The R package phyloseq[26] (v. 1.20.0) was used for storing the ASV table, taxonomy, and associated sample data and for calculating alpha diversity measures.

### Contaminant identification and removal

Decontam[20] (v.0.8) was used to identify ASVs with significant inverse correlations with DNA concentration measured by Nanodrop (prior to library preparation). We evaluated the performance of decontam using different user-defined thresholds.

SourceTracker[21] version 1.0.1 was used to determine the amount of contamination in samples and recover the true microbial community of each sample. To test optimal inputs as source environments, we performed a series of analyses with the following samples labeled as source samples: contaminant profiles (created by removing the expected ASVs from the diluted mock microbial community samples), blank control (negative extraction controls that went through all the sample processing steps and microbial free water), and the expected mock microbial community profile (the undiluted mock microbial community sample).

The data and R markdown scripts necessary to reproduce the analysis presented in this paper is available at https://github.com/lakarstens/ControllingContaminants16S.

## Results

The total number of paired-end reads from the mock community dilution samples was 2,555,160. Following quality filtering, a total of 1,325,836 sequences were further analyzed and grouped into 755 amplicon sequence variants (ASVs). Each sample in the mock community dilution series had between 36,014–223,112 reads. The undiluted mock community sample (D0) had 222,159 reads. As expected, the number of reads per sample generally decreased with increased dilution, though not linearly (Figure 1A, Table 1). The first round of dilution (D1) had 152,456 reads, and the following 3 rounds of dilution (D2-4) had between 194,418 and 223,112 reads per sample. D5 and D6 had between 113,844 and 124,657 reads per sample, whereas D7 and D8 had 36,014 and 38,405 reads per sample, respectively. The blank control sample had 182,624 reads (Table 1).

**Table 1.**
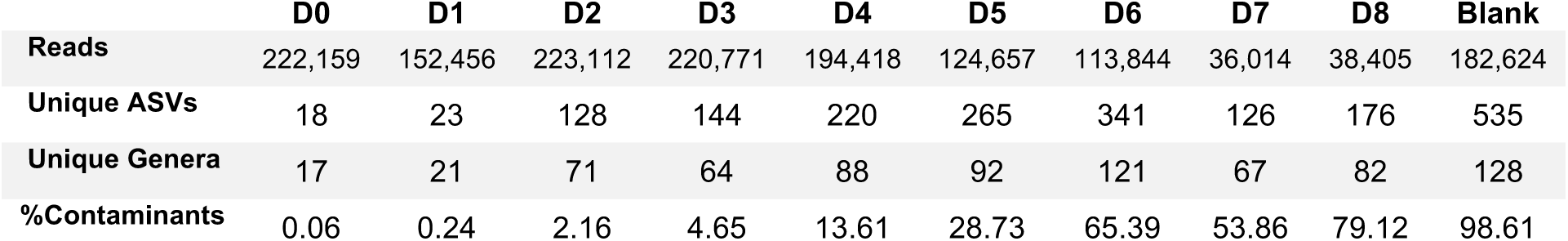
The impact of decreasing starting material for 16S rRNA gene sequencing.

|  | D0 | D1 | D2 | D3 | D4 | D5 | D6 | D7 | D8 | Blank |
| --- | --- | --- | --- | --- | --- | --- | --- | --- | --- | --- |
| Reads | 222,159 | 152,456 | 223,112 | 220,771 | 194,418 | 124,657 | 113,844 | 36,014 | 38,405 | 182,624 |
| Unique ASVs | 18 | 23 | 128 | 144 | 220 | 265 | 341 | 126 | 176 | 535 |
| Unique Genera | 17 | 21 | 71 | 64 | 88 | 92 | 121 | 67 | 82 | 128 |
| %Contaminants | 0.06 | 0.24 | 2.16 | 4.65 | 13.61 | 28.73 | 65.39 | 53.86 | 79.12 | 98.61 |

**Figure 1.**
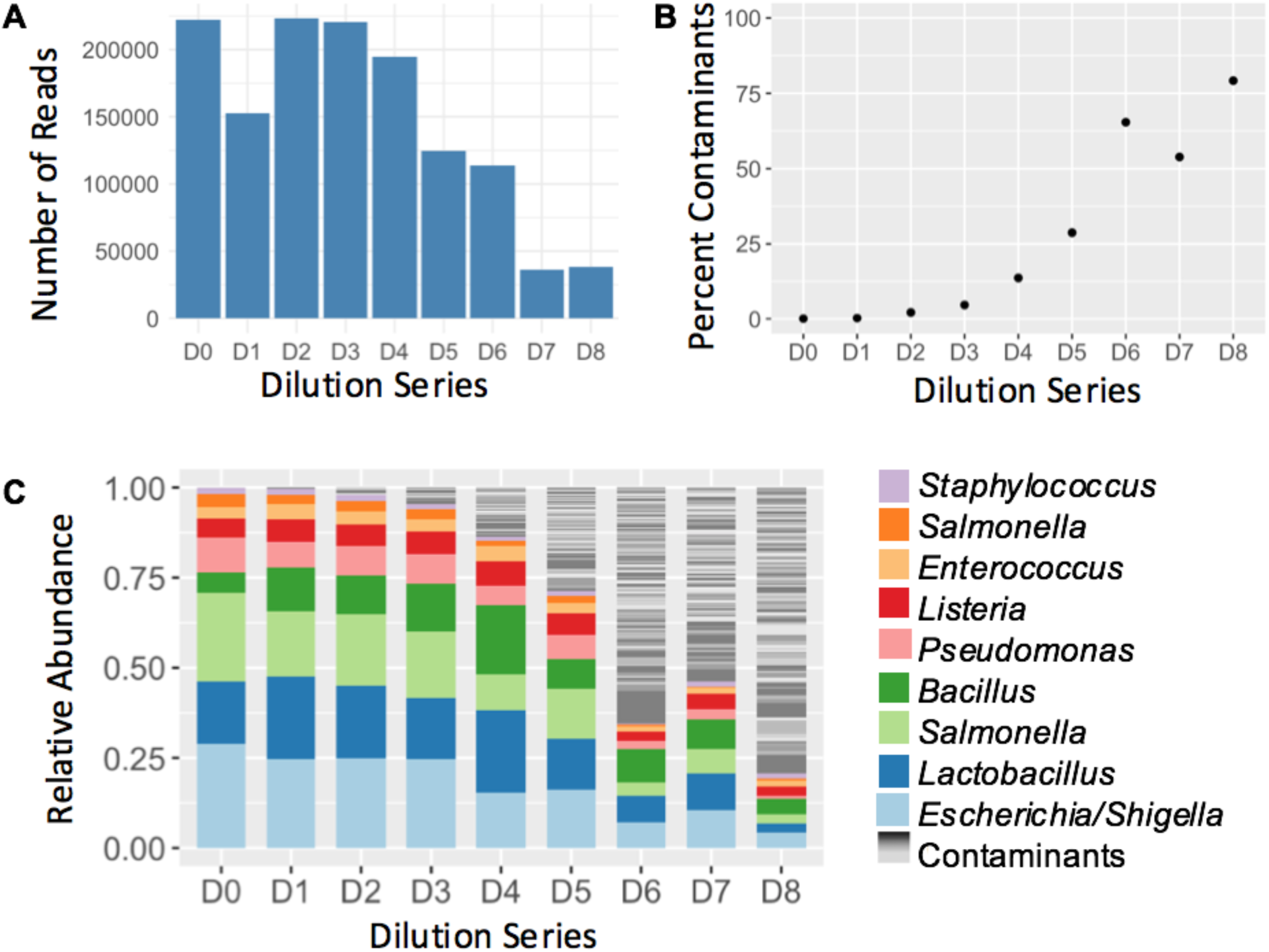
Analysis of a mock microbial community dilution series reveals contaminating bacteria increases with decreasing starting DNA. A mock microbial community consisting of eight known bacteria was subjected to eight rounds of a threefold dilution (1:3 through 1:6561), followed by bacterial DNA isolation and amplification, and sequencing with Illumina MiSeq. **A.** The number of reads per sample varied between 223,112 and 36,014 reads. **B.** The proportion of reads from contaminant DNA increased with the amount of dilution. **C.** Stacked bar plot representing the bacteria identified in each sample. The expected sequences from the mock microbial community are displayed in color; all other bacterial sequences are in grayscale.

### Undiluted mock community sample composition

The undiluted mock community contained 18 unique ASVs, which corresponded to 17 unique genera. The expected mock community genera were represented in the top 9 abundant ASVs. One expected genus, *Salmonella*, was split into two ASVs based on a single nucleotide variation (G to T transition). The other 9 unexpected ASVs totaled 0.05% of reads; they were not similar to the expected sequences in the mock community nor were they identified as chimeric sequences based on the sequence composition of the mock community. Due to their low relative abundance and lack of other explanation, these were likely either contaminants from sample processing or barcode crosstalk from sequencing.

### Blank extraction control composition

The blank control was composed of 535 ASVs that mapped to 128 unique genera. The most abundant genera was *Bacteroides* (12.9%) followed by *Ruminiclostridium_6* (6.4%), and *Prevotellaceae_UCG-001* (6.0%). Twenty-three ASVs were present at abundances between 1% and 5%, and the remaining 116 ASVs were present at relative abundances less than 1%. 5 of the 8 mock community genera were present in the blank control. These were in low abundance (0.02% to 1.9%), making up 3.7% of the total relative abundance.

### Mock community dilution series composition

The amount of contaminant DNA increased with subsequent dilutions of the mock microbial dilution series, indicated by an increased number of ASVs and bacterial genera (Table 1) that were not in the expected 16S rRNA sequences from the mock microbial community. As expected, the number of ASVs and number of genera increased with decreased starting microbial biomass for D1-D6. However, this trend did not hold for the most diluted samples (D7 and D8), which had 126 and 176 ASVs corresponding to 59 and 73 genera (Table 1). This could be due to the lower number of reads from these samples. We did identify an increased percentage of reads not originating from the expected mock community sequences, which had a relationship with dilution (Figure 1B).

The mock microbial community was composed of 8 bacteria that resulted in 9 unique ASVs. Thus, there were 746 ASVs in the dilution series that were not the expected mock microbial community ASVs and are referred to as **contaminant ASVs**. After two rounds of dilution, contaminant ASVs became more predominant in the microbial community profiles (Figure 1B, 1C), with contaminants having relative abundances greater than 1% of sequence reads. After the 6^th^ round of dilution (D6), contaminants contribute over 50% to the estimated community composition. Importantly, contaminant genera in these dilutions were detected in relative abundances greater than 10%, making individual contaminants easily mistaken for true microbial composition of a sample.

The contaminants identified with abundances greater than 1% are listed in Supplemental Table 2. The most prevalent ASV was *Bacteroides* (species unclassified), present with relative abundances up to 9.1% in the most dilute samples (D6-8), followed by two ASVs identified as *Lachnospiraceae NK4A136* group with relative abundances up to 3.7%. Other contaminant ASVs had less than 2% relative abundance per sample. 696 of the contaminant ASVs had a relative abundance greater than 0.01% in at least one sample.

199 of the 746 contaminant ASVs were also present in the blank control sample (see Supplemental Table 3). ASVs from the blank extraction control made up 37.98%-66.25% of the contaminant sequences per sample (average 51.57%). 547 ASVs were unique to the dilution series and not present in the blank control. Of these, 493 ASVs were only present in a single dilution series sample, with 2–485 reads per sample (mean 76.89). Contaminant ASVs that were not present in the blank control made up less than 8% of the ASVs per sample.

### Impact of contaminants on alpha diversity metrics

To identify the impact of including contaminants in downstream microbiome analyses, we calculated commonly used measures of alpha diversity: observed number of ASVs, the Shannon index, and the Inverse Simpson index. The expected values for these measures based on the expected mock microbial community sequences are 9, 1.85, and 5.20, respectively. The inclusion of contaminant sequences led to increases in all estimates (Chao1 estimator 18–341, Shannon index 1.85–4.69, Inverse Simpson index 5.20–67.78 Figure 2).

**Figure 2.**
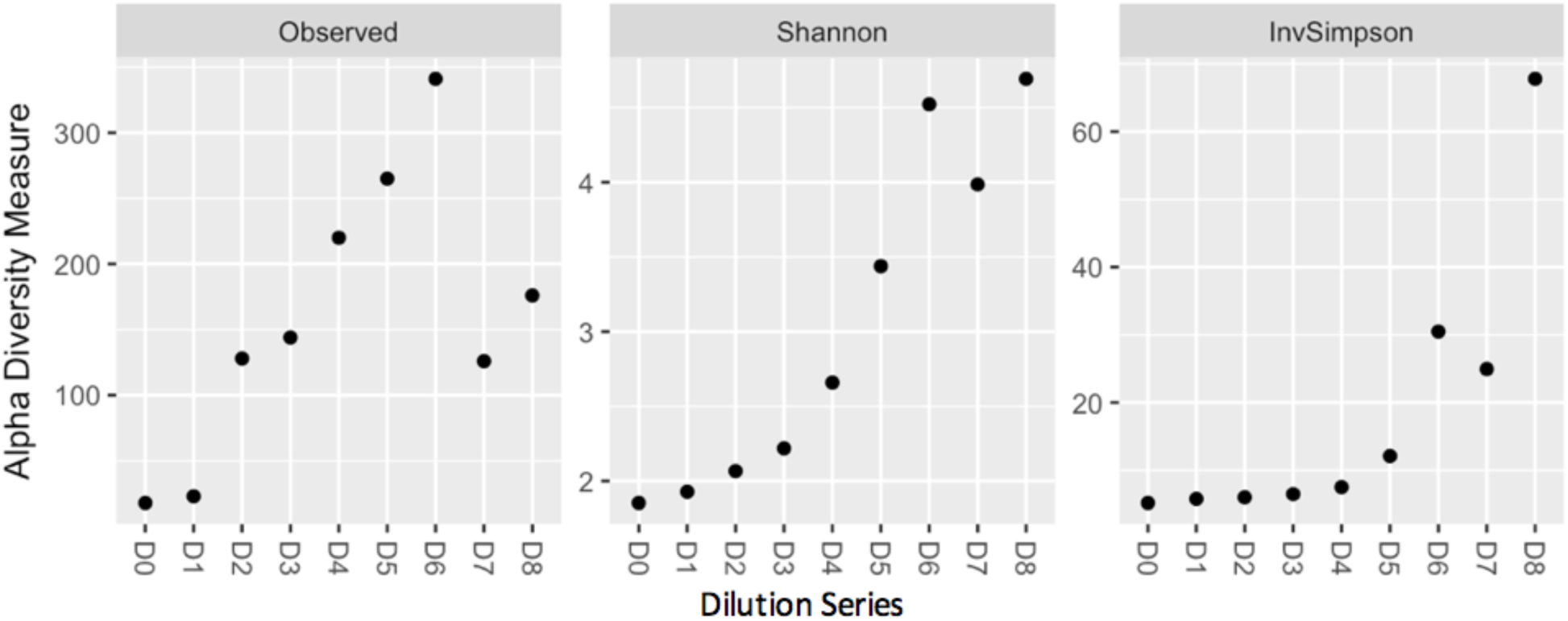
Impact of contaminants on common alpha diversity measures. Failure to remove contaminants from the dataset leads to increase in observed number of ASVs and estimates of alpha diversity evaluated by the Shannon diversity index and the Inverse Simpson index.

### Contaminant removal by correlation with DNA concentration with decontam

An inverse relationship between amplicon concentration after library preparation and number of reads has been reported for both Illumina sequencing[9] and pyrosequencing[15]. This provides an objective way to identify contaminants, and has been implemented in the open-source R package decontam[20] through the ‘frequency’ method. We evaluated decontam with the frequency method using a variety of thresholds, ranging from the default 0.1 to 0.5. As expected, increasing the threshold lead to an increased proportion of ASVs being identified as contaminants across the dilution series (Table 2). Importantly, decontam did not remove any expected mock microbial community sequences.

**Table 2.**
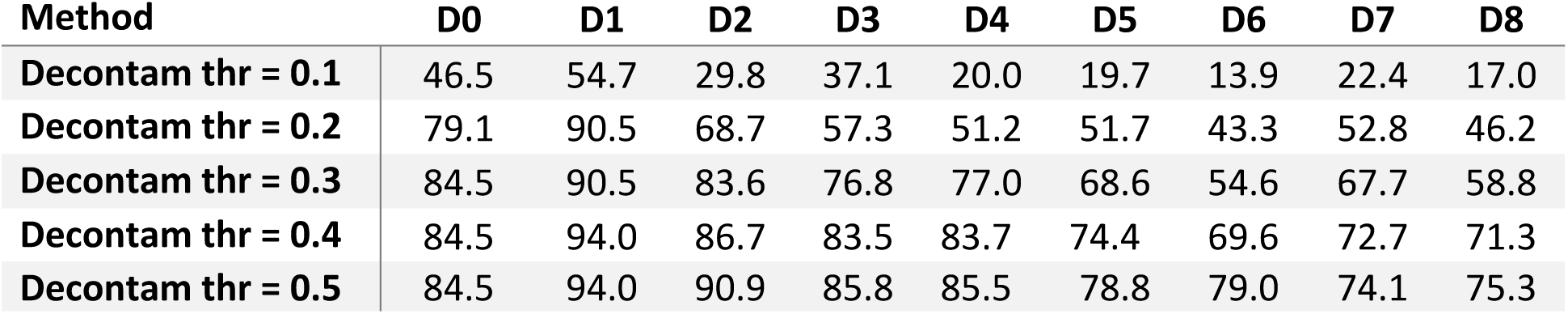
Percent of contaminants removed with decontam at varying thresholds (thr) 0.1–0.5.

Using decontam with a threshold of 0.5, 240 ASVs were identified as contaminants across all samples (Figure 3), which made up 74.1–94% of the contaminants per sample (Table 2). As depicted in Figure 3C, decontam removed the majority of contaminants in D1-5, though contaminant ASVs still made up more than 20% of the more dilute samples (D6-8). Additional filtering steps such as low abundance filtering or the prevalence method available in decontam would likely further remove these contaminant ASVs.

**Figure 3.**
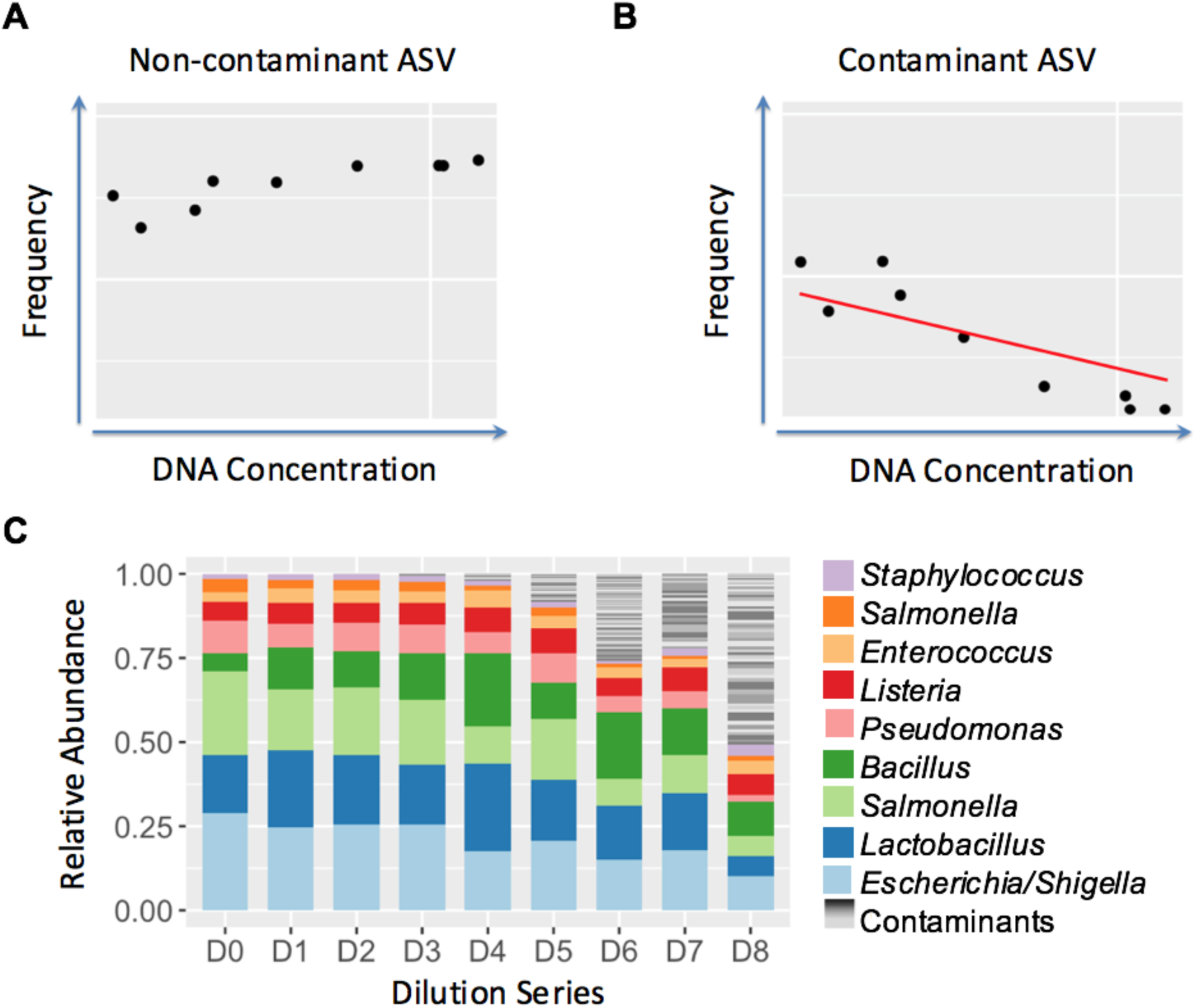
Removal of contaminant ASVs with the frequency method in the R package decontam. **A.** An example of a non-contaminant ASV. The abundance of this ASV does not have a relationship with DNA concentration. **B.** An example of a contaminant ASV detected by decontam. The abundance of the ASV has a negative relationship with DNA concentrations. **C.** The mock microbial dilution series with contaminants removed. Decontam did not remove any of the expected mock microbial community sequences and removed the majority of contaminants. In highly dilute samples (D6–D8), some contaminant sequences are still present in the sample (making up between 23–48% of the corrected microbial profile), but is reduced from the original data (**see** Figure 1).

### Contaminant removal with SourceTracker

We used the SourceTracker algorithm[21] to identify and remove contaminant sequences from the mock microbial dilution series. SourceTracker predicts the proportion of unknown ‘sink’ samples that arise from defined ‘source’ samples. We tested two scenarios for recovering the expected mock microbial community profiles from the mock microbial dilution series. In the first scenario, the expected mock community profile from the undiluted mock microbial community sample served as a source environment, mimicking the scenario when the experimental source environment is well defined. In the second scenario, the expected mock microbial community is unknown; the proportion of sequences not predicted to be from the defined blank control or contaminant profile sources is called the contamination-corrected profile. The second scenario is the more commonly encountered scenario, where the low microbial biomass environment that is being studied is poorly defined. For each scenario, we additionally evaluated the use of a combination of blank control and contaminant profiles (Case 1) or the blank control alone (Case 2) as source environments.

### Scenario one: Well-defined experimental source environment

The first scenario is one in which the experimental source environment is well defined, with the undiluted mock microbial community sample (D0) defined as a source environment. The contaminant source environments were defined as both contaminant profile and blank control (Case 1). To test if the blank controls alone were enough to identify contaminants, we additionally used SourceTracker with D0 and the blank control as source environments (Case 2).

SourceTracker was able to remove the majority of contaminants, with less than 0.01% contaminants remaining in the most dilute samples (Figure 4A and 4B). However, SourceTracker also removed some reads of mock microbial community taxa, though this removal was minimal (< 6.2 %). With both contaminant profiles and blank controls defined as sources, SourceTracker identified the majority of reads as arising from contaminant profiles and less than 0.1% classified as blank or unknown, and less than 1% of the mock microbial reads misclassified (Figure 5B). With the blank control only defined as a contaminant source environment, the majority of reads arising from contaminant sources were classified as unknown, with a maximum of 4.2% of reads being classified as arising from the blank control (Figure 5C). Additionally, a greater proportion of the mock microbial community sequences were classified incorrectly as unknown (maximum 6.2%).

**Figure 4.**
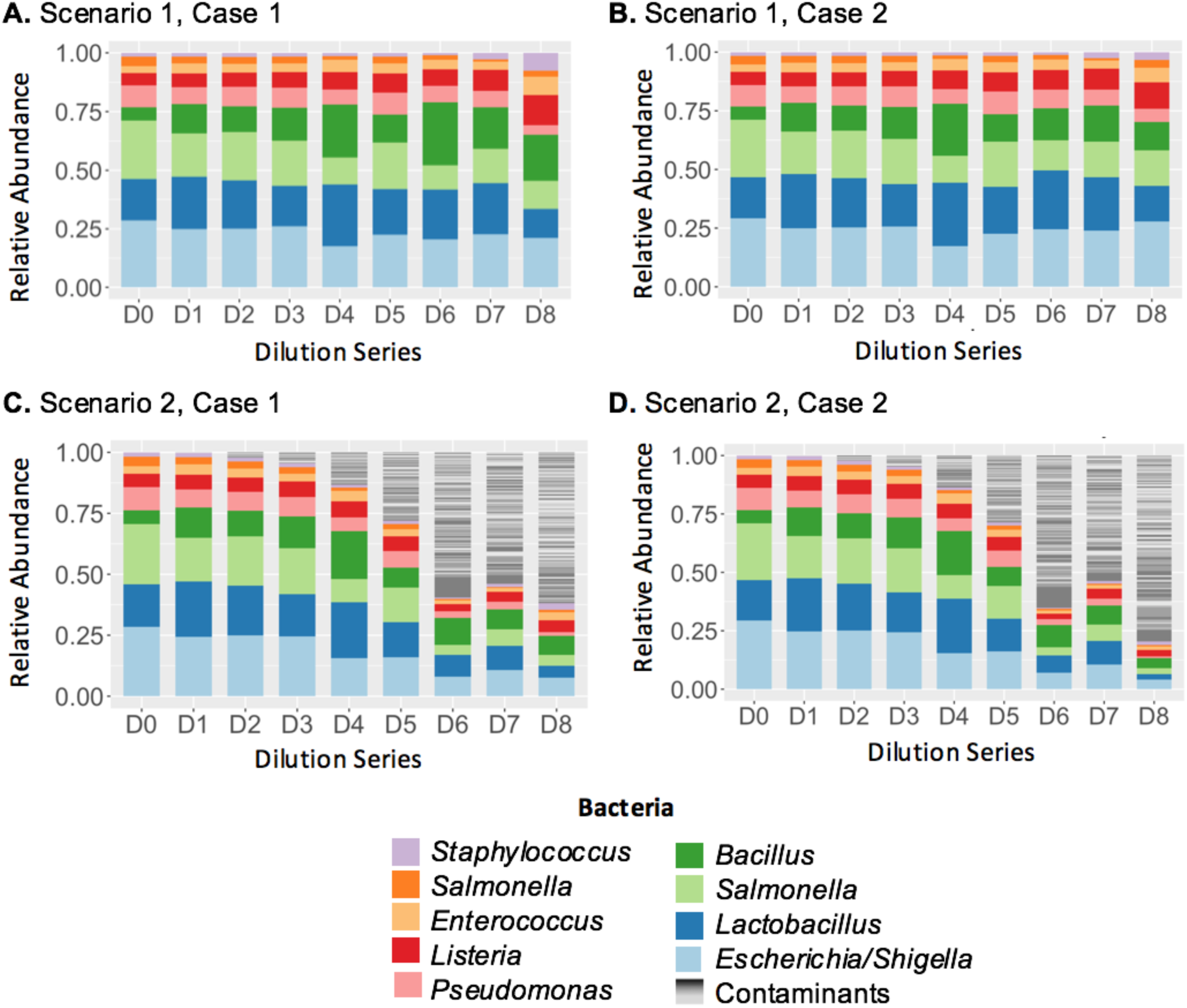
Removal of contaminants with SourceTracker. **A.** Scenario 1, Case 1: Recovered mock microbial community profiles from the scenario where both the experimental and contaminant environments are well defined. In this scenario, SourceTracker performs excellently. **B.** Scenario 1, Case 2: Recovered mock microbial community profiles from the scenario where the experimental environment is well defined and only the blank control is defined as a contaminant source environment. **C.** Scenario 2, Case 1: Recovered mock microbial community profiles from the scenario where the contaminant environments are defined as contaminant profiles and blank control and the experimental environments are not defined. The recovered profiles still contain the majority of contaminant sequences (gray). **D.** Scenario 2, Case 2: Recovered mock microbial community profiles from the scenario where the only defined environment is a contaminant environment defined by a blank control.

**Figure 5.**
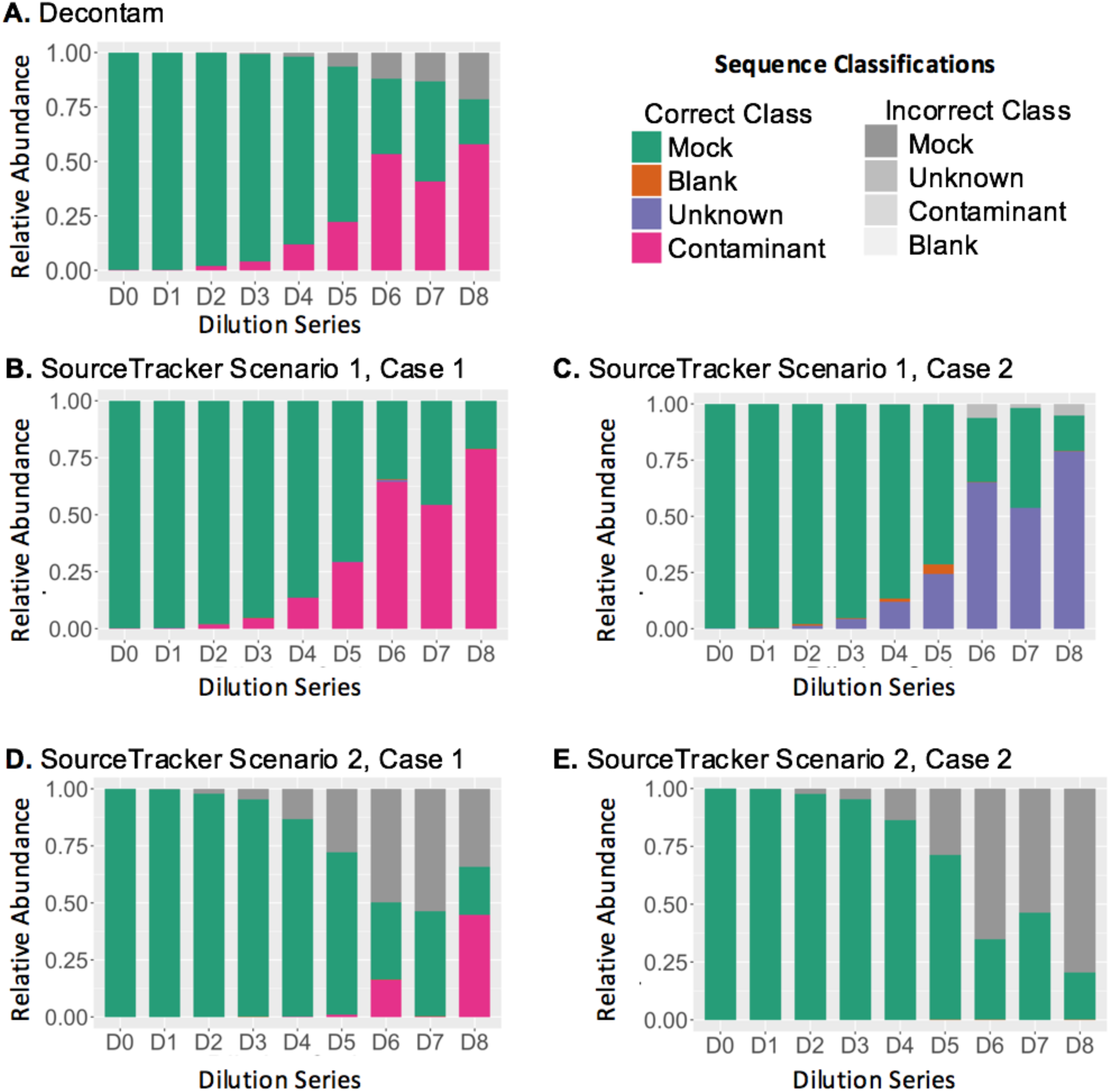
Summary of contaminant removal performance. **A.** Results from using decontam to classify sequences as contaminant or non-contaminant. The majority of sequences are classified correctly, though the proportion of contaminants misclassified as non-contaminants (grey mock) increases with increasing dilution. **B.** Results from the SourceTracker scenario where both the experimental and contaminant environments are well defined. The samples are primarily classified correctly. Some of the expected mock microbial community reads were classified as Unknown (maximum 0.014). **C.** SourceTracker results from the scenario where the experimental environment is well defined and only the blank control is defined as a contaminant source environment. SourceTracker performs well in this scenario in recovering the mock microbial community; however, the majority of contaminants are classified as Unknown rather than Blank, and a greater proportion of the mock microbial community sequences are classified incorrectly as Unknown. **D.** SourceTracker results from the scenario where only contaminant environments are defined as contaminant profiles and blank control and the experimental environment is unknown. In most samples, less than 1% of reads were classified as arising from the contaminant profile or blank profile, with the exception of D6 and D8 where a greater proportion of reads were correctly identified as contaminants. **E.** SourceTracker results from the scenario where the contaminant environment is defined by a blank control and the experimental environment is unknown. The majority of contaminant reads are classified incorrectly and less than 1% of reads per sample were correctly classified as arising from the blank source environment.

### Scenario two: Unknown experimental source environment

To mimic the common experimental scenario where the expected source environment is not well defined, we used SourceTracker with only the contaminant and blank profiles defined as the source environments. In this design, any bacterium that is classified as ‘unknown’ by SourceTracker would be assumed to come from the ‘unknown’ environment of interest and thus represent the contaminant-corrected microbial profile. In this scenario, SourceTracker failed to identify most contaminants, leaving 0.06–68.9% of contaminants (Figure 4C and Figure 4D).

### Overall performance of contaminant removal methods

To evaluate the overall performance of each contaminant removal method, ASVs were classified as being correctly or incorrectly identified as mock community, blank control, or contaminants (Figure 5). Decontam did not misclassify any mock community sequences as contaminants and correctly classified the majority of contaminant sequences correctly as contaminants (74 – 91%, Figure 5A). SourceTracker with well-defined experimental environments (Scenario One) also classified the majority of ASVs correctly (94 – 100%), though the proportion of contaminants misclassified as non-contaminants increased with increasing dilution (Figure 5B and 5C). SourceTracker with only contaminant environments defined and an unknown experimental environment (Scenario two) did not correctly classify many (0 – 56%) of the contaminant ASVs (Figure 5D and 5E).

## Discussion

### Impact of contaminants on microbial community composition

Our study demonstrates the impact of laboratory contaminants on 16S rRNA gene sequencing experiments of varying starting microbial biomass on a mixed microbial community. The diluted mock microbial community samples showed a marked increase in the number of ASVs and thus genera with decreasing starting material. This led to increased estimates of commonly used alpha diversity metrics incorporating species or genus richness. Highly dilute samples also demonstrated contaminant bacteria that were higher in abundance than the expected sequences. The inclusion of such large amounts of contaminant taxa can artificially decrease the relative abundance of the actual expected microbial communities. Our findings demonstrate the magnitude of error from biological datasets harboring contaminants unaddressed by suitable control procedures.

Contaminated laboratory reagents in 16S rRNA gene based experiments has long been recognized in the scientific literature[27]. Several groups have recently investigated the impact of bacteria arising from DNA extraction kits and other commonly used laboratory reagents specifically on 16S rRNA gene sequencing experiments. Salter et al. performed a serial dilution of a pure bacterial isolate to assess contaminants[28]. Similar to our study, after 4 ten-fold dilutions of a single bacterial isolate, they identified that samples were predominantly composed of contaminant DNA sequences. Glassing et al. demonstrated the effect of contaminants on biological samples of low microbial biomass and evaluated methods for mitigating contaminant contributions using a variety of negative controls[10]. These studies exemplify the need for researchers to be adamant about contaminant control procedures used.

### Identifying contaminants in low microbial biomass experiments

Using a dilution series of mock microbial communities, where the expected composition and sequences of the mock microbial community are known, we also evaluated two available analytical methods to minimize and remove contaminant sequences from downstream analyses. The use of mock microbial communities as a positive control for 16S rRNA gene sequencing experiments has been advocated[29], though it is unclear how often they are used in practice. Mock microbial communities are composed of mixtures of known bacterial composition and are subject to the same experimental and computational processing as experimental samples. Since the expected composition of the mock microbial community is known, they can be used to identify problems in the experimental protocol, to understand biases introduced in the experimental protocol (i.e. PCR amplification bias), to optimize the bioinformatics workflow used, or to develop new methods.

Our data suggests that the use of negative controls alone are likely insufficient to inform researchers of appropriate measures to minimize contaminants from their experiments. While we found that sequences present in the blank control sample were also present in the mock community dilutions, we also identified many unique sequences as contaminants in the mock microbial dilution samples that were absent from the blank control. Furthermore, we found that five out of eight of the expected sequences from the mock microbial community were also present in the blank extraction control. Removing all sequences present in the blank extraction controls would have led to erroneously removing the majority of the expected mock community dilution sequences. Glassing et al. also demonstrated that contaminant removal by removing sequences present in the blank control was too harsh. Thus, using mock microbial dilutions in studies of low microbial biomass environments as controls may provide a more accurate method to evaluate and determine contaminant sequences than negative controls alone.

### Removing contaminants in low microbial biomass experiments

Analytical and bioinformatics approaches have been suggested as a post-processing step to identify and remove potential contaminants, but to our knowledge this is the first study examining the performance of these methods at varying microbial biomass. These methods include identifying correlations with OTU / ASV frequency and starting DNA material measured either before library preparation with Pico-Green or Nanodrop[9] or by more accurate but more labor-intensive qPCR[19]. We evaluated the former as implemented in the decontam package available in R[20], which demonstrated reasonable performance for removing the majority of contaminants, though some contaminants still remained in the most dilute samples. Another approach uses Bayesian modeling to estimate the proportion of contaminants as implemented in SourceTracker.

Our study indicates that using SourceTracker is an excellent approach if the environment under investigation is well defined, removing 99% of contaminants. However, SourceTracker performed poorly in the scenario where the environment under investigation is unknown, which is likely for most low microbial biomass environments. In this scenario, SourceTracker removed less than 1% of contamiants for most samples, and removed 56% for one sample. Decontam performed well, removing 100–75% of contaminants, and did not depend on prior knowledge about the microbial community being studied.

Another consideration for selecting a contaminant removal method is the format of the results, which may limit downstream analysis. Decontam classifies OTUs / ASVs as either being contaminants or not, and OTUs / ASVs that are identified as contaminants are removed from the entire dataset. This has the benefit of retaining the raw read count of the data, but does not permit removal of a proportion of a contaminating sequence, which may be desirable if a known contaminant is also known to be part of the experimental environment being studied. SourceTracker has the benefit of estimating proportions of sequences that arise from a contaminant source rather than removing the entire sequence from the dataset. However, SourceTracker takes a significant time to run, which increases with sequencing depth. To overcome this, it is common/recommended practice to subsample the data prior to using SourceTracker, which is currently a controversial practice in microbiome analysis and limits the downstream analyses that can be performed[30].

### Limitations

Our study used a commercial mock microbial community consisting of 8 known bacteria and 2 fungi. Mock microbial communities that are representative or have similar characteristics to the expected experimental microbial community can also be generated and may provide a more robust estimate of the experimental contaminants present in a study. Furthermore, in our study, all blank controls were barcoded with the same barcode, limiting their utility. For example, we were unable to use the prevalence method for contaminant removal that is available in the decontam package.

## Conclusions

Controlling for contaminants in low microbial biomass experiments remains an important and unsolved problem, particularly in the age of easily accessible high throughput next generation sequencing. We demonstrate that using a dilution series of a mock microbial community can provide researchers studying low microbial biomass environments an effective means to evaluate laboratory contaminants in 16S rRNA gene sequencing experiments. This includes identifying the real background noise specific for a particular experiment, evaluating effective strategies and appropriate parameters to remove contaminant DNA sequences, and a means to evaluate the success of methods used to identify exogenous contaminant DNA from experiments. Since the exogenous DNA present in DNA extraction kits have been shown to vary across kit types and different lots of the same kit, a mock community dilution series is an important addition to each 16S rRNA gene sequencing experiment in order to identify laboratory-and experiment-specific contaminants. This is particularly important for studies investigating environments with low microbial biomass such as urine, placenta, and lung.

Our recommendations are for investigators to use a mock community dilution series consisting of at least a high–medium–and low dilution representing the expected range of DNA concentration in the environment being studied in addition to negative extraction blanks as negative controls. We also strongly recommend that newer bioinformatics pipelines using methods based on amplicon sequence variants (ASVs) rather than operational taxonomic units (OTU) are used since the former approach offers finer resolution, which is particularly important for distinguishing contaminant sequences. Importantly, it is likely that one single approach for removing contaminants from low biomass samples will NOT work equivalently across all sequencing studies investigating low microbial biomass environments. Thus, it is important that the method used for controlling contaminants be monitored and reported upon publication. This will ensure that science is transparent and will make results more robust and reproducible.

## Availability of data and materials

The sequencing data supporting the conclusions of this article is available in the SRA under accession number: PENDING.

## Author Contributions

Conceived and designed experiments: LK and MA. Performed experiments: LK, MA, SD. Analyzed the data: LK. Wrote the paper: LK. Advised on interpretation of data: SKM, JB, DF, TG, MA. Critical revision of manuscript: WTG, AJW, JB, SKM.

## Acknowledgements

We thank the researchers who developed DADA2, phyloseq, decontam, and SourceTracker for making their programs open source and available to the scientific community.

This research was supported by the following: LK was supported as a Scholar of the Oregon BIRCWH K12 Program funded by the Eunice Kennedy Shriver National Institute of Child Health & Human Development of the National Institutes of Health (NIH) under Award Number K12HD04348. JB was supported by NIH UL1TR001881 and DK46763. AJW was supported by NIH R01 DK104718. SKM was supported by NIH/NCATS UL1TR002369. LK and MA were supported by the MA was supported by the Rheumatology Research Foundation. MA was also supported by the Spondylitis Association of America

The content is solely the responsibility of the authors and does not necessarily represent the official views of the National Institutes of Health

---

**Supplemental Table 1.** Estimated number of bacterial cells that yielded the DNA used in 16S rRNA gene PCR reaction for the mock microbial dilution series

| Samples | Concentration of Samples<br>(cells/ml) | Total bacteria used for<br>extraction | Estimated number of<br>bacterial cells that yielded the<br>DNA used in 16S rRNA gene<br>PCR reaction |
| --- | --- | --- | --- |
| D0 | 1.40E+10 | 1.05E+09 | 21,000,000 |
| D1 | 4.67E+09 | 3.50E+08 | 7,000,000 |
| D2 | 1.56E+09 | 1.17E+08 | 2,333,333 |
| D3 | 5.19E+08 | 3.89E+07 | 777,778 |
| D4 | 1.73E+08 | 1.30E+07 | 259,259 |
| D5 | 5.76E+07 | 4.32E+06 | 86,420 |
| D6 | 1.92E+07 | 1.44E+06 | 28,807 |
| D7 | 6.40E+06 | 4.80E+05 | 9,602 |
| D8 | 2.13E+06 | 1.60E+05 | 3,201 |

**Supplemental Table 2.**
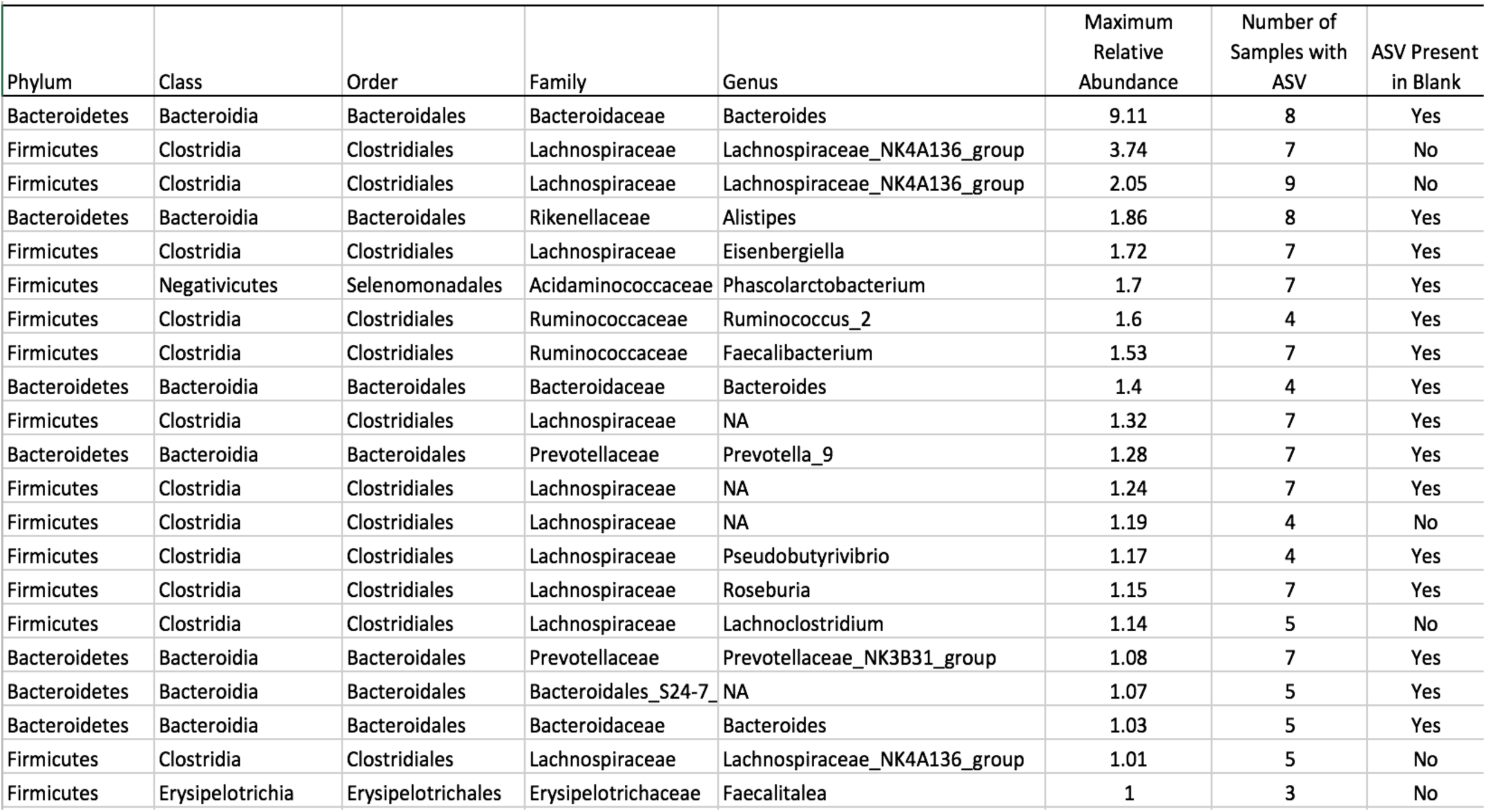
Taxanomic classification and sequence of contaminantswith atleaset 1% relative abundance in atleaset one sample.

**Supplemental Table 3.** Summary of contaminant sequences across the dilution series

| Measure | Number of ASVs |
| --- | --- |
| Unexpected ASVs | 746 |
| Present in all samples | 1 |
| Present in at least 3 samples | 153 |
| Present in 1 sample | 493 |
| Present in Blank | 199 |
| Relative abundance >0.01% in at least 1 sample | 724 |

## References

1. Dickson RP, Erb-Downward JR, Martinez FJ, Huffnagle GB. The Microbiome and the Respiratory Tract. Annu Rev Physiol. 2015; October 2015:1–24. DOI:10.1146/annurev-physiol-021115-105238.

2. Aagaard K, Ma J, Antony KM, Ganu R, Petrosino J, Versalovic J. The placenta harbors a unique microbiome. Sci Transl Med. 2014;6:237ra65. DOI:10.1126/scitranslmed.3008599.

3. Karstens L, Asquith M, Davin S, Stauffer P, Fair D, Gregory WT, et al. Does the urinary microbiome play a role in urgency urinary incontinence and its severity? Front Cell Infect Microbiol. 2016;6 JUL:1–13. DOI:10.3389/fcimb.2016.00078.

4. Brubaker L, Wolfe AJ. The new world of the urinary microbiota in women. Am J Obstet Gynecol. 2015;213:644–9. DOI:10.1016/j.ajog.2015.05.032.

5. Pearce MM, Zilliox MJ, Rosenfeld AB, Thomas-White KJ, Richter HE, Nager CW, et al. The female urinary microbiome in urgency urinary incontinence. Am J Obstet Gynecol. 2015;213:347.e1-11. DOI:10.1016/j.ajog.2015.07.009.

6. Tanner MA, Goebel BM, Dojka MA, Pace NR. Specific Ribosomal DNA Sequences from Diverse Environmental Settings Correlate with Experimental Contaminants. Appl Environ Microbiol. 1998;64:3110–3.

7. Salter S, Cox MJ, Turek EM, Calus ST, Cookson WO, Moffatt MF, et al. Reagent contamination can critically impact sequence-based microbiome BMC. analyses Biol. 2014;:7187. DOI:10.1101/007187.

8. Lusk RW. Diverse and widespread contamination evident in the unmapped depths of high throughput sequencing data. PLoS One. 2014;9.

9. Jervis-Bardy J, Leong LEX, Marri S, Smith RJ, Choo JM, Smith-Vaughan HC, et al. Deriving accurate microbiota profiles from human samples with low bacterial content through post-sequencing processing of Illumina MiSeq data. Microbiome. 2015;3:19. DOI:10.1186/s40168-015-0083-8.

10. Glassing A, Dowd SSE, Galandiuk S, Davis B, Chiodini RJRRJ, Pace N, et al. Inherent bacterial DNA contamination of extraction and sequencing reagents may affect interpretation of microbiota in low bacterial biomass samples. Gut Pathog. 2016;8:24. DOI:10.1186/s13099-016-0103-7.

11. Weiss S, Amir A, Hyde ER, Metcalf JL, Song SJ, Knight R. Tracking down the sources of experimental contamination in microbiome studies. Genome Biol. 2014;15:564. DOI:10.1186/s13059-014-0564-2.

12. Champlot S, Berthelot C, Pruvost M, Bennett EA, Grange T, Geigl EM. An efficient multistrategy DNA decontamination procedure of PCR reagents for hypersensitive PCR applications. PLoS One. 2010;5. DOI:10.1371/journal.pone.0013042.

13. D’Amore R, Ijaz UZ, Schirmer M, Kenny JG, Gregory R, Darby AC, et al. A comprehensive benchmarking study of protocols and sequencing platforms for 16S rRNA community profiling. BMC Genomics. 2016;17:55. DOI:10.1186/s12864-015-2194-9.

14. Wright ES, Vetsigian KH. Quality filtering of Illumina index reads mitigates sample cross-talk. BMC Genomics. 2016;17:876.

15. Willner D, Daly J, Whiley D, Grimwood K, Wainwright CE, Hugenholtz P. Comparison of DNA Extraction Methods for Microbial Community Profiling with an Application to Pediatric Bronchoalveolar Lavage Samples. PLoS One. 2012;7:e34605. https://doi.org/10.1371/journal.pone.0034605.

16. Bittinger K, Charlson ES, Loy E, Shirley DJ, Haas AR, Laughlin A, et al. Improved characterization of medically relevant fungi in the human respiratory tract using next-generation sequencing. Genome Biol. 2014;15:487.

17. Flores R, Shi J, Yu G, Ma B, Ravel J, Goedert JJ, et al. Collection media and delayed freezing effects on microbial composition of human stool. Microbiome. 2015;3:33.

18. Adams RI, Bateman AC, Bik HM, Meadow JF. Microbiota of the indoor environment: a meta-analysis. Microbiome. 2015;3:49.

19. Lazarevic V, Gaïa N, Girard M, Schrenzel J. Decontamination of 16S rRNA gene amplicon sequence datasets based on bacterial load assessment by qPCR. BMC Microbiol. 2016;16:73. DOI:10.1186/s12866-016-0689-4.

20. Davis NM, Proctor D, Holmes SP, Relman DA, Callahan BJ. Simple statistical identification and removal of contaminant sequences in marker-gene and metagenomics data. bioRxiv. 2017. DOI:10.1101/221499.

21. Knights D, Kuczynski J, Charlson ES, Zaneveld J, Mozer MC, Collman RG, et al. Bayesian community-wide culture-independent microbial source tracking. Nat Methods. 2011;8:761–3. DOI:10.1038/nmeth.1650.

22. Callahan BJ, Mcmurdie PJ, Rosen MJ, Han AW, Johnson AJ, Holmes SP. DADA2: High resolution sample inference from amplicon data. Nat Method. 2016;13 August 2015:581–3. DOI:10.1101/024034.

23. Callahan BJ, McMurdie PJ, Holmes SP. Exact sequence variants should replace operational taxonomic units in marker-gene data analysis. ISME J. 2017;11:2639–43. http://dx.doi.org/10.1038/ismej.2017.119.

24. Tikhonov M, Leach RW, Wingreen NS. Interpreting 16S metagenomic data without clustering to achieve sub-OTU resolution. ISME J. 2015;9:68–80.

25. Wang Q, Garrity GM, Tiedje JM, Cole JR. Naive Bayesian classifier for rapid assignment of rRNA sequences into the new bacterial taxonomy. Appl Environ Microbiol. 2007;73:5261–7. DOI:10.1128/AEM.00062-07.

26. McMurdie PJ, Holmes S. Phyloseq: An R package for reproducible interactive analysis and graphics of microbiome census data. PLoS One. 2013;8.

27. Corless CE, Guiver M, Borrow R, Kaczmarski EB, Fox AJ. Contamination and sensitivity issues with a Real-Time Universal 16S rRNA PCR. 2000;38:1747–52.

28. Salter SJ, Cox MJ, Turek EM, Calus ST, Cookson WO, Moffatt MF, et al. Reagent and laboratory contamination can critically impact sequence-based microbiome analyses. BMC Biol. 2014;12:87. DOI:10.1186/s12915-014-0087-z.

29. Bokulich NA, Rideout JR, Mercurio WG, Shiffer A, Wolfe B, Maurice CF, et al. mockrobiota: a public resource for microbiome bioinformatics benchmarking. mSystems. 2016;1.

30. McMurdie PJ, Holmes S. Waste Not, Want Not: Why rarefying microbiome data is inadmissible. PLOS Comput Biol. 2014;10:e1003531. https://doi.org/10.1371/journal.pcbi.1003531.

